# iPSC-Derived Cerebral Organoids Reveal Mitochondrial, Inflammatory and Neuronal Vulnerabilities in Bipolar Disorder

**DOI:** 10.1101/2025.02.25.640229

**Authors:** Dana El Soufi El Sabbagh, Alencar Kolinski Machado, Lauren Pappis, Erika Leigh Beroncal, Delphine Ji, George Nader, Prathyusha Ravi Chander, Jaehyoung Choi, Angela Duong, Hyunjin Jeong, Bruna Panizzutti, Chiara Cristina Bortolasci, Andrea Szatmari, Peter Carlen, Margaret Hahn, Liliana Attisano, Michael Berk, Ken Walder, Ana Cristina Andreazza

## Abstract

Bipolar disorder (BD) is increasingly recognized as a disorder with both mitochondrial dysfunction and heightened inflammatory reactivity, yet their contribution to neuronal activity remains unclear. To address these gaps, this study utilizes iPSC-derived cerebral organoids (COs) from BD patients and healthy controls to model disease-specific metabolic and inflammatory dysfunction in a physiologically relevant system. BD COs exhibited mitochondrial impairment, dysregulated metabolic function, and increased nod-leucine rich repeat and pyrin domain containing protein 3 (NLRP3) inflammasome activation sensitivity. Treatment with MCC950, a selective NLRP3 inhibitor, effectively rescued mitochondrial function and reduced inflammatory activation in both BD and control COs. Additionally, a Bioactive Flavonoid Extract (BFE) was explored as a potential therapeutic, demonstrating partial rescue of inflammasome activation. These findings highlight a mitochondria-inflammasome axis in BD pathophysiology and establish a novel platform for studying BD-associated cellular mechanisms, ultimately bridging the gap between molecular dysfunction and therapeutic development.

## Introduction

Bipolar Disorder (BD) is a complex mood disorder characterized by recurrent episodes of mania, hypomania, and depression, affecting approximately 1% of the global population, with onset typically in late adolescence or early adulthood ^1^. Early diagnosis is associated with better prognosis; however, delays of approximately nine years from the initial symptom onset remain common ^1^. Long-term treatment relies on mood stabilizers, antipsychotics, and antidepressants, yet over 50% of patients experience inadequate symptom control, contributing to chronic morbidity^1^. In addition to psychiatric symptoms, BD is associated with higher prevalence of metabolic syndrome, cardiovascular disease (CVD), and obesity, leading to a reduction in life expectancy by 12–14 years, primarily due to premature cardiovascular mortality^2^. Both genetic and environmental factors, including childhood trauma, infections, lifestyle factors (e.g., sleep disturbances, physical inactivity), and substance use, contribute to disease susceptibility^2^. Neuroimaging studies have consistently reported structural abnormalities in the prefrontal cortex, along with reduced neuronal density in BD patients^3^ ^4^, emphasizing the need to better understand the underlying biological mechanisms of the disorder.

Healthy mitochondrial function is essential to maintain the brain’s high energy demands as the brain utilizes 20% of whole-body energy. Mitochondria also regulate crucial pathways including calcium homeostasis and reactive oxygen species (ROS) production which are critical for neuronal plasticity^5^. Bioenergetic and mitochondrial dysfunction in psychiatric disease has been studied as early as 1926, and subsequent research has further confirmed the crucial role of this organelle^6^. Post-mortem brains and peripheral samples from BD patients have repeatedly shown altered mitochondrial genetics encoding for electron transport chain (ETC) complexes, oxidative stress imbalances, as well as increased levels of circulating cell free mitochondrial DNA (ccf-mtDNA)^7–10^. At the cellular level, emerging evidence suggests a strong link between mitochondrial dysfunction and inflammation in BD pathophysiology. Mitochondria play a crucial role in cellular homeostasis; however, dysfunction can lead to a multifactorial phenotype involving mtDNA release, dysregulated bioenergetics, and altered neurogenesis as the mitochondria fail to meet the cells’ energy demands ^11,12^.

Patients with BD are also known to have heightened levels of pro-inflammatory cytokines both systemically and in the CSF^13^. We have previously reported in a cohort of 837 patients with BD that ccf-mtDNA can act as a marker of chronic low-grade inflammation, a hallmark of BD^2^. Chronic inflammation has long been implicated in BD with increased levels of pro-inflammatory cytokines especially in manic episodes^13^. ccf-mtDNA can directly promote inflammation by acting as a damage-associated molecular pattern (DAMP) which can activate numerous pathways to heighten the immune response. These include the cGAS-STING pathway which increases interferons (IFNs), the NF-kB pathway which upregulates nod-leucine rich repeat and pyrin domain containing protein 3 (NLRP3), and binding to toll-like receptors (TLRs) to activate the NLRP3 inflammasome^14,15^. Based on previous work from our group, increased NLRP3 inflammasome activation has found in pre-frontal post-mortem brain samples and in peripheral blood mononuclear cells (PBMCs) of patients with BD^14^. NLRP3 has been repeatedly proposed as a link between mitochondrial dysfunction and neuroinflammation, as dysfunctional mitochondria produce high levels of ROS that can act as a DAMP and are recognized by pattern recognition receptors (PRRs) which activate the inflammasome ^16,17^. Further, mitochondria can become permeabilized because of NLRP3 activation^15^. All these processes contribute to altered mitochondrial bioenergetics by decreasing the activity of the ETC, altering adenosine triphosphate (ATP) levels, depolarizing mitochondrial membrane potential (MMP), and increasing ROS damage^15,18^.

This bi-directional relationship between mitochondrial dysfunction and inflammation in BD warrants further investigation, as these factors can have serious consequences on overall brain health when mitochondria fail to re-establish intra and extracellular homeostasis, further impacting BD severity (Figure 1). Due to limitations in modelling BD, patient-derived induced pluripotent stem cells (iPSCs) offer a unique and transformative approach. These cells enable the generation of cerebral organoids (COs) to study this disorder and its underlying cellular and mitochondrial mechanisms in a physiologically relevant environment. COs recapitulate the 3-dimensiolnal structure and cellular diversity of a fetal brain, allowing for an in-depth exploration into the cellular interactions of BD-specific phenotypes^19^. Using COs provides a powerful platform to investigate the interplay between mitochondrial dysfunction and inflammation in BD, bridging the gaps in our understanding of the disease.

**Figure 1.**
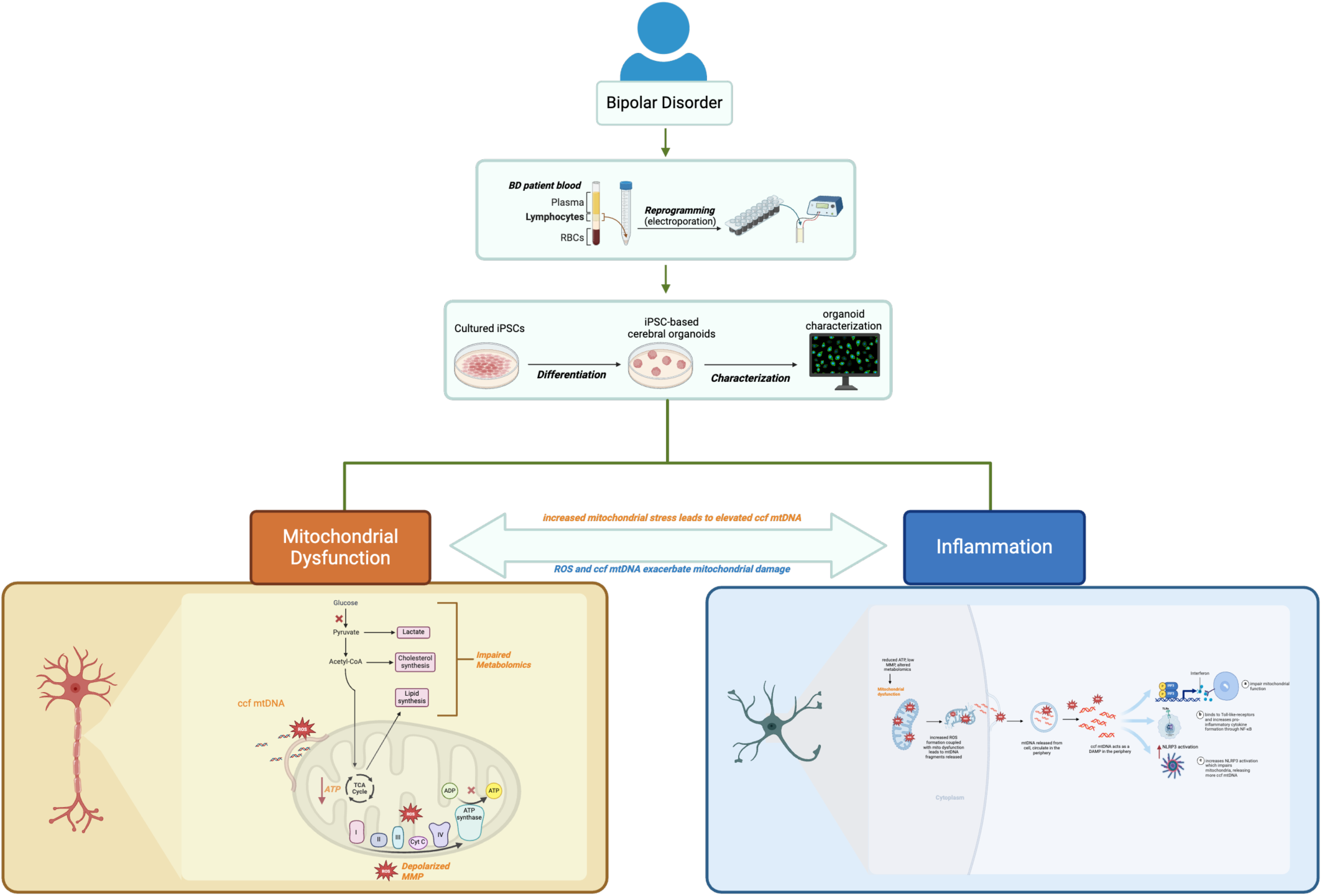
Overview of study design. 3 BD and 3CT patient blood was collected and PBMCs were isolated and reprogrammed by electroporation into iPSCs. iPSCs were cultured and characterized then COs were generated and they underwent numerous assays to investigate the role of mitochondrial dysfunction and inflammation in BD.

To address these gaps, this study utilizes iPSC-derived COs from individuals with BD and healthy controls to model disease-specific mitochondrial and inflammatory dysfunction in a physiologically relevant system. By leveraging this advanced 3D model, we investigate mitochondrial health, key metabolic vulnerabilities, neuronal activity changes, and inflammatory susceptibilities associated with BD. We hypothesize that BD-derived COs will exhibit mitochondrial dysfunction and heightened inflammatory reactivity, which contribute to abnormal neuronal firing. Furthermore, we explored COs as a tool to validate potential therapeutics, specifically, the potential of a Bioactive Flavonoid Extract (BFE), to mitigate NLRP3 inflammasome activation and restore cellular homeostasis. This study provides a novel platform to dissect BD-associated cellular mechanisms and identify potential therapeutic targets, bridging the gap between molecular findings and clinical interventions.

## Methods

### Study design

This study began with the recruitment of five deeply-phenotyped patients diagnosed by a psychiatrist with BD (BDI or BDII) and five age- and sex-matched non-psychiatric controls. These individuals underwent comprehensive clinical and biomarker assessments before their PBMCs were isolated for iPSC reprogramming. Following iPSC characterization, only three BD-derived and three control-derived iPSC lines met quality control criteria and were further used to generate COs.

The COs were employed as a patient-specific 3D model to investigate mitochondrial health, cellular stress responses, and inflammatory activation sensitivity in BD pathology. Mitochondrial health was assessed through metabolomic profiling, bioenergetic analysis, and structural imaging, while inflammation was evaluated via NLRP3 inflammasome activation sensitivity. Additionally, COs were treated with a leading molecule that blocks the NLRP3 activation and with BFE, a novel therapeutic to explore potential anti-inflammatory rescue effects. This multi-faceted approach provides a comprehensive framework for understanding the cellular and mitochondrial mechanisms underlying BD, offering insights into neuroinflammation, mitochondria metabolic dysregulation, and therapeutic modulation.

### Blood Sample Collection and Processing

This study was conducted in partnership with the IMPACT Institute at Deakin University (Victoria, Australia). We followed the guidelines implemented by the Biomarkers Task Force as listed by the World Federation of Societies of Biological Psychiatry for blood sample collection, clinical assessment, and ethics^20^. This study was performed and approved by the Research Ethics Board at the University of Toronto, Ontario, Canada using REB (36359) and at Deakin University with REB (17/205) in accordance with the Helsinki Declaration of 1975. Dr. Andreazza’s laboratory has also received approval from the Stem Cell Oversight Committee (Canadian Institutes of Health Research, #399222).

Five adults diagnosed with bipolar disorder (BDI or BDII) were recruited from IMPACT Institute and Barwon Health, Australia by a licenced psychiatrist (Dr. Michael Berk) using the Structured Clinical Interview for Diagnostic and Statistical Manual 5 (DSM-5 Disorders) (SCID-5). Five non-psychiatric healthy controls were recruited in the Greater Geelong Area by local advertisements. Clinical demographics were collected and informed consent of participants was obtained. Venous blood was drawn (30mL) into lavender-top EDTA coated tubes (Vacutainer Plus).

Peripheral blood was diluted 1:2 in wash buffer (PBS containing 2% FBS and 2 mM EDTA), then carefully layered onto SepMate™-15 tubes (StemCell Technologies, Canada) prefilled with Lymphoprep (StemCell Technologies, Canada). The samples were centrifuged at 1200g for 10 minutes to separate the PBMCs. The PBMC layer was collected, washed with the same buffer, and centrifuged at 300 g for 10 minutes. To eliminate residual red blood cells, the pellet was briefly rinsed with ice-cold Milli-Q water before another wash with buffer, followed by a final centrifugation at 300 g for 10 minutes. PBMCS were cryopreserved in FBS +10% DMSO with slow-freezing containers in -80°C and moved to liquid nitrogen the following day. For plasma, 1 EDTA tube (different from the one used for PBMCs) was centrifuged for 10 minutes at 4°C and 1,300g. Resulting plasma was aliquot into low-binding cryotubes and stored at -80°C until analysis. Samples were shipped to the University of Toronto under dry ice and PBMCs were thawed and washed twice with DPBS and 2×10^6^ cells were seeded in a tissue culture flask in 10mL of StemPro-34 SFM (10639011 Gibco) with GlutaMAX (1X), recombinant human GM-CSF(25ng/mL), recombinant human IL-3 (50ng/mL), and recombinant human SCF (100ng/mL) for 72 h at 37 °C with 5% CO_2_ and humidity.

### Generation and characterization of induced pluripotent stem cells from PBMCs

Details of iPSC generation and characterization are described in Duong et al (2021)^21^. In short, PBMCs were transfected using Epi5 EBNA-1/OriP episomal vectors containing 5 pluripotent factors (OCT4, SOX2, LIN28, KLF4, and L-MYC). Electroporation occurred using he Lonza 4D-Nucleofactor System (Lonza, AAF-1002B). Cells were then transferred onto a 6 well plate coated with Geltrex (Gibco). IPSCs were grown in mTesR media (StemCell Technologies) and characterized for pluripotency using an EpiPluriScore (Cygenia) and immunofluorescence. IPSCs also underwent karyotyping analysis to confirm successful reprogramming. Successful karyotyping enabled continuation of the study and in this case, 3 out of 5 BD and 3 out of 5 controls moved forward with the study.

### Cerebral organoid (CO) generation

COs were generated from patient iPSCs. In summary, iPSCs were singularized and plated in 96well V bottom plates (Greiner) and allowed to form embryonic bodies (EBs). After a few days of neural induction, the EBs were embedded in Matrigel (Corning) to induce a 3D shape. The COs were then cut out of Matrigel and placed on an orbital shaker to grow into 3D uniform round structures. More details on the methods of generation can be found in Sivitilli et al. (2020)^21–23^.

### Singularizing and counting cells of COs

To singularize COs, first, COs were first washed twice with PBS then 2 mL of Accutase^TM^ (AT-104, Innovative Cell Technology Inc., San Diego, CA, USA) was added. Steps are described in detail in Duong et al for single-cell dissociation of COs^21,22^. The singularized cells were counted on Orflow (MXF001) and resuspended in the required assay-specific buffer, or frozen as cell pellets for metabolomic profiling.

### Metabolomics

Both patient plasma and intracellular CO metabolomic profiling was conducted in collaboration with the University of Ottawa’s Metabolomics Core Facility. For plasma, the plasma was frozen at -80°C and sent on dry ice to the facility. For COs, the metabolomics were quantified from cells using liquid chromatography mass spectrometry (LC-MS) for over 20 mitochondrial-related metabolites. All the samples analysed were conducted by a blinded analysis. More details on metabolite selection and processing can be found in Zachos et al. (2024)^2^.

### Intracellular ATP measurements

Intracellular ATP levels were measured using Cell Titer Glo Luminescent Cell Viability Assay (Promega, G7570) according to manufacturer’s instructions. COs were singularized and cells were seed in a density of 50,000 cells/well per 100μL Hanks’ Balanced Salt Solution (HBSS) buffer in a 96-well white polystyrene plate (Greiner CELLSTAR, 655083). ATP standard curve (0nM-10μM) was generated using ATP disodium salt (Sigma-Aldreich, A7699). Luminescence was measured using a Synergy H1 microplate reader equipped with Gen 5 software (BioTek Instruments, Inc., 253147)^23^.

### Immunofluorescence microscopy

The CO slices were fixed with 4% of paraformaldehyde (PFA) overnight at 4°C, and the next day washed with PBS then placed in 30%(w/v) sucrose for another overnight incubation at 4°C. The samples were freeze embedded on OCT (Tissue-Tek) and cryosectioned on a 20 μM thickness for immunofluorescence stain. Cryosectioned slices were stained as previously described by Duong et al (2021)^21^ and El Sophie El Sabbagh (2024)^22^ using MAP2 for neurons (I3-1500) (1:200), SOX2 for neural progenitors (MAB2018) (1:200), GFAP for astrocytes (Abcam, Cambridge, UK, #ab4674) (1:500) and ASC for ASC specks (AL177) (1:200) antibodies to assess the NLRP3 inflammasome assembly and ASC specks formation on the astrocytes. Secondary antibodies used were Donkey anti-Rabbit IgG [H + L], secondary antibody Alexa Fluor Plus 647 (Invitrogen #A32795), Goat anti-mouse [IgG] [H + L] (Abcam, #Ab97935) secondary antibody Cy3, and Donkey anti-Chicken IgY [H + L] secondary antibody Alexa Fluor 488 (Jackson ImmunoResearch Laboratories Inc., West Grove, PA, USA, #703-546-155). The antibodies were diluted with 0.5% BSA, and after staining, coverslips were mounted with ProLong Gold Antifade Mountant containing DAPI (Invitrogen, Waltham, MA, USA, P36935), and imaging of the samples was performed at the Microscopy Imaging Laboratory (MIL), University of Toronto, Canada, using an LSM 880 Elyra Superresolution confocal microscope (Zeiss). After staining, images were quantified for area DAPI, area SOX2, area GFAP, and area MAP2. Cells that were double positive for GFAP and SOX2 were referred to as GFAP progenitors. This analysis was performed using an automated image analysis software HALO Indica Labs (v4.0.5107.357) using the analysis tool Area Quantification F (v4.2.3).

### Transmission Electron Microscopy (TEM)

Mitochondrial morphology and number were evaluated using transmission electron microscopy (TEM) in the University of Toronto’s MIL. The CO’s were fixed with a primary fixation buffer (1% glutaraldehyde in 0.1M phosphate buffer and 4% paraformaldehyde) for 2h at room temperature and changed to fresh fixation buffer for an overnight incubation at 4°C. After the incubation, samples were washed with 0.1 M phosphate buffer 3 times for 20 min. Following the washes, samples were incubated for 1h at room temperature with a second fixing buffer (1% osmium tetroxide in phosphate buffer). The following steps were performed at the microscope imaging laboratory, University of Toronto (MIL): Dehydration was conducted with graded series of ethanol (50%; 70%, 90% and 100%). After dehydration the CO were washed with propylene oxide twice for 15min and infiltrated with a graded series of a mixture of epoxy resin and propylene oxide following Duong ate al (2021)^21^. For the polymerization step, COs were transferred to a Been embedding capsule and polymerized at 60°C for 48h. The resulting resin block was sectioned to a thickness of 90 nm using a Reichert Ultracut E microtome. Each section was stained with saturated uranyl acetate for 15 minutes, washed with distilled water, and then incubated for an additional 15 minutes with Reynold’s lead citrate. The samples were washed again with water and subsequently prepared for microscopy using an FEI Talos L120C transmission electron microscope at an accelerating voltage of 80 kV^21^.

### CO acute slice preparation

COs were washed twice with PBS, then placed in base molds containing 4% low melting-point agarose in PBS on ice for 10 min. After the samples froze, COs were sliced using a vibratome (Leica Biosystems VT1200) on a 400 μm thickness. The slice was removed from the vibratome and placed in a 6 well plate with PBS and 1% v/v Penicillin-Streptomycin (PenStrep) (Gibco 15070). The plate was moved to a Biological Safety Cabinet and the sliced CO was transferred to a new 6 well plate containing CO media with PenStrep^21,22^.

### Mito Tracker staining

The CO slices were prepared as described in acute slice preparation and placed in media containing Mito Tracker Red CMX Ros (Invitrogen, M7512) at 100 nM for 30 min at 30°C in the dark. Other slices were incubated with Mito Tracker Green (MitoTracker Green, Invitrogen). After the incubation, samples were washed twice with PBS for 5 minutes each on the shaker and then taken for live imaging at the Microscopy Imaging Laboratory (MIL), University of Toronto, Canada, using an LSM 880 Elyra Superresolution confocal microscope (Zeiss).

After staining, images were quantified for area DAPI and area mitotracker red or green using an automated image analysis software HALO Indica Labs (v4.0.5107.357) using the analysis tool Area Quantification F (v4.2.3).

### Assessment of Mitochondria Membrane Potential (MMP)

CO slices were incubated with JC-1 ((5,5′,6,6′-tetrachloro-1, 1′,3,3′-tetraethylbenzimi-dazolylcarbocyanine iodide) Dye (Invitrogen, Mitochondrial Membrane Potential Probe, T3168) at a concentration of 1 μg/mL for 30 min in the dark on an orbital shaker at 37°C. After the incubation, samples were washed three times with PBS, and DAPI (Invitrogen) was added on the second wash. Immediately after the washes, regular media was added to the CO slices and live imaging was performed at the MIL, University of Toronto, Canada, using an LSM 880 Elyra Superresolution confocal microscope (Zeiss). After staining, images were quantified for area DAPI and area red and area green using an automated image analysis software HALO Indica Labs (v4.0.5107.357) using the analysis tool Area Quantification F (v4.2.3).

### Electrophysiology recordings

Each incubated CO slice was placed in a recording chamber and perfused with artificial cerebrospinal fluid (ACSF) [125 mM NaCl, 25 mM NaHCO3, 2.5 mM KCl, 10 mM Glucose, 2.5 mM CaCl2, 1.5 mM MgCl2], bubbled with carbogen (95% O2, 5% CO2) and maintained at a temperature of 35-37°C. Local field potentials (LFPs) were recorded using borosilicate capillary glass electrodes (6–10 MΩ) filled with ACSF. The electrodes were positioned away from the necrotic center and approximately 100 µm deep in the healthy peripheral tissue. Electrode placement was verified using an upright Olympus BX51 WI fluorescent microscope equipped with a 4× water immersion objective lens (Olympus Corporation). Signals were acquired with a Digidata 1322A digitizer (Axon Instruments) and a Multiclamp 700A amplifier (Molecular Devices), using PClamp software (version 10.7) at a sampling rate of 25,000 Hz. LFPs were continuously recorded for approximately 15-30 minutes from all slices. Physiological data were analyzed using custom-written software in a MATLAB environment. Following each day’s experiments, the slices were discarded.

### Extracellular markers assays

#### Circulating cell free mitochondria DNA (ccf-mtDNA)

DNA extraction was performed using QiaAMP DNA mini kit (Qiagen) from cell culture supernatant. After the DNA extraction, ccf-mtDNA was quantified using Taqman^TM^ Duplex polymerase chain reaction (PCR) with primers and probe targeting β2M and PPIA for nuclear DNA, and ND1 and ND4 for mitochondrial DNA (Table 2). PCR was performed on a CFX96 Real Time Thermocycler (BioRad), using Commercial oligonucleotide in a concentration ranging from 108 to 102 copies/µL as a standard curve. Thermocycler conditions were 50 °C for 2 min, 95 °C for 20 s, followed by 40 cycles of 95 °C for 3 s, and 60 °C for 30 s. Results were expressed as ccf-mtDNA copies/µL^22,24^. Refer to Table 1 for sequences.

**Table 1.**
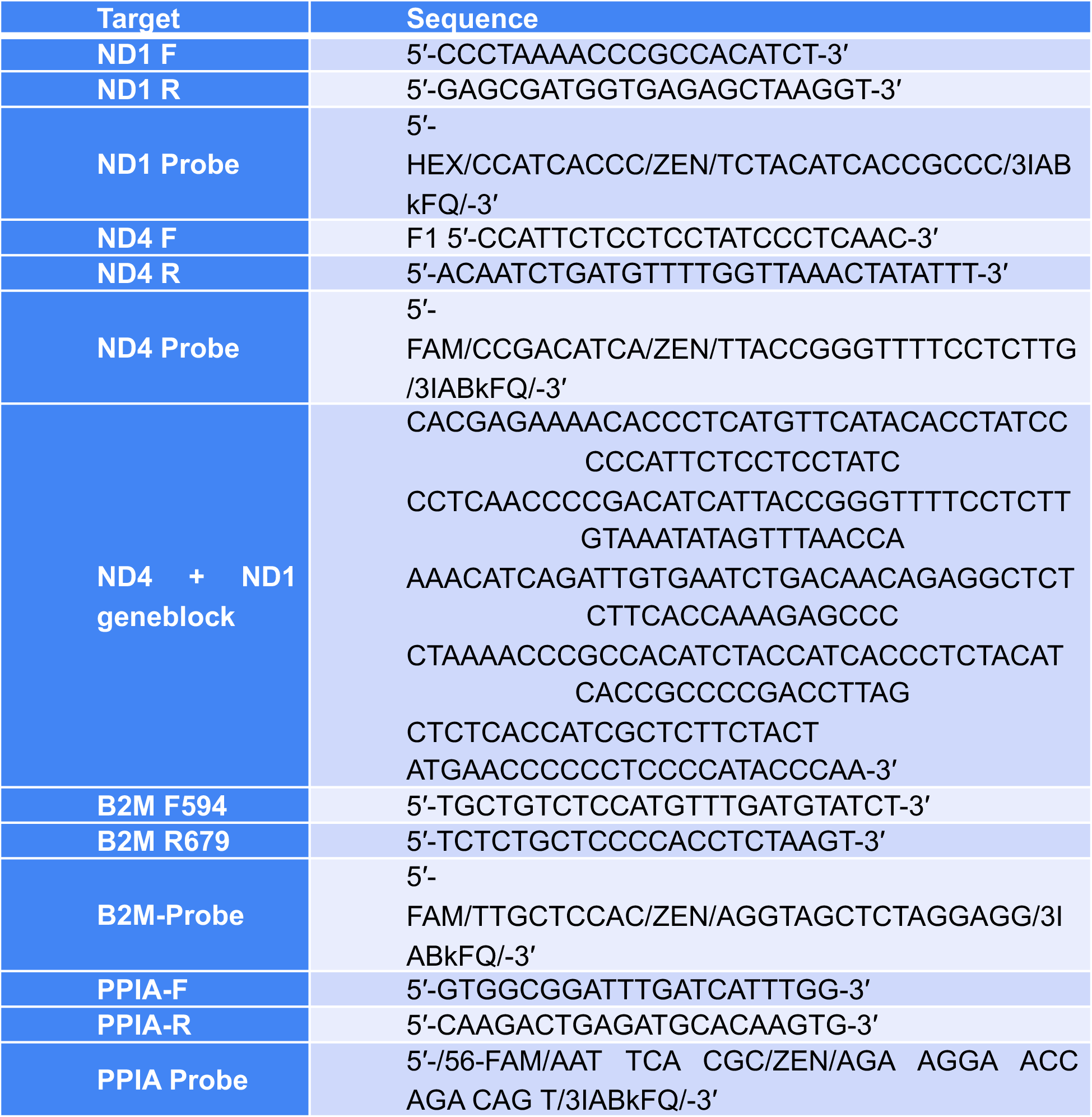
Probes and primers sequences for ccf-mtDNA PCR.

**Table 2.** Epi-Pluri-Score analysis of iPSC lines.

| Sample ID | DNA Methylation percentage at 3 CpGs |  |  | Epi-Pluri-Score |
| --- | --- | --- | --- | --- |
| | β-value<br>[ANKRD46] | β-value<br>[C14orf115] | β-value<br>[POU5F1] | |
| CT002 | 18.9 | 9.8 | 75.2 | 9.1 |
| CT003 | 26.9 | 6.7 | 59.1 | 20.2 |
| CT004 | 26.1 | 8.9 | 77.4 | 17.1 |
| BD001 | 23.9 | 6.7 | 59 | 17.2 |
| BD002 | 21.7 | 9.9 | 70.1 | 11.8 |
| BD003 | 16 | 9.6 | 72.2 | 6.4 |

### Quantification of Reactive Oxygen Species (ROS) production

The total level of ROS production was measured by using 2′,7′-dichlorodihydrofuorescein diacetate (DCFH-DA) (Sigma-AldrichD6883) following instructions by Degli et al (2009)^25^. Fluorescence intensity was measured using a Synergy H1 microplate reader equipped with Gen 5 software (BioTek Instruments, Inc., Winooski, VT, USA, 253147). Drug treatment effects were calculated by percentage compared to the untreated group.

### Quantification of dsDNA release

The extracellular release of dsDNA was assessed via DNA PicoGreen fluorescent probe (Quant-iT^TM^ PicoGreen^TM^, Thermo Fisher-P11495; Eugene, OR, USA) following Ahn et al. (1996)^26^. Fluorescence was recorded at 480 nm excitation and 520 nm emission using a Synergy H1 microplate reader equipped with Gen 5 software (BioTek Instruments, Inc., Winooski, VT, USA, 253147). Drug treatment effects were calculated by percentage compared to the untreated group.

### Activation of NLRP3 inflammasome

NLRP3 inflammasome activation was conducted according to the methods described in El Soufi El Sabbagh et al. (2024)^22^. Briefly, CO slices were treated with lipopolysaccharide (LPS) (Invivogen, San Diego, CA, USA, TLRL-3PELPS LPS) at 100 ng/mL for 3h as a priming step. Next, the LPS was removed and 10 μM of nigericin (Sigma, St. Louis, MO, USA, N7143) was added for a 4-hour incubation as the activator step of the NLRP3 inflammasome.

### Preparation of the Bioactive Flavonoid Extract (BFE)

The BFE is based on a well-known Brazilian super antioxidant berry called *Euterpe oleracea* Mart., (acai). The development, characterization and matrix composition of this extract can be found in detail in our previous publications^27–29^. The BFE safety profile was characterized in previous published studies using neuron-like cells, microglia cell lines, THP-1 drived macrophages, A549 lung cells and lung organoids^30^.

### Inhibiting the NLRP3 inflammasome

For inhibition of the inflammasome, 100nM of MCC950, a leading molecule for NLRP3 inflammasome inhibition (Invivogen, San Diego, CA, USA, 210826-40-7) was used as a 2-hour pretreatment between the priming (LPS) and activating (nigericin) steps. To assess BFE’s efficacy in comparison to MCC950, BFE at 1 μg/mL (optimal concentration stablished and published by Machado et al., 2016; Souza et al., 2020; and Cadona et al., 2021, ADD LUNG PAPER) for 2h, also in between the LPS and nigericin exposures to evaluate its potential on inhibiting the NLRP3 protein multicomplex activation in sliced COs^27,29,31^. At the end of all treatments, the CO slices were fixed and stained for immunofluorescence imaging and the supernatant was collected for analyses of extracellular markers.

### Quantification of ASC specks and NLRP3 Activation and Rescue

The quantification of ASC specks was performed using methods described in El Soufi El Sabbagh et al. (2024)^22^. To count the ASC specks a semi-automated method was employed using FIJI software (ImageJ, version 2.9.0/1.53t), that recognized the brightness, size and shape of the ASC specks. Each image was then assigned a random number to ensure anonymity, and all detected specks were independently counted by a blinded observer. Each image also went through automated cell count in HALO Indica Labs (v4.0.5107.357) using the analysis tool Cytonuclear FL (v4.2.3) which counts and characterizes cells in an automated manner.

To assess NLRP3 activation, values were normalized to baseline conditions for both CT and BD COs. Activation (LPS+Nigericin treatment) was expressed as a percentage of baseline, where baseline values were set to 100%, and post-stimulation values were calculated as:

% Activation = (Activated Condition-Baseline Condition)×100.

To quantify differences in activation sensitivity, the fold difference between BD and CT was calculated using: fold difference = (BD activation %)/ (CT activation %). A secondary percentage activation bar graph was generated to illustrate the relative increase in activation across conditions, emphasizing the greater activation response in BD COs compared to CT. Statistical comparisons were performed using an unpaired t-test or two-way ANOVA with post-hoc corrections, as appropriate.

To quantify the rescue effect of MCC950 or BFE treatment on NLRP3 activation, the percentage rescue was calculated relative to the activated positive control (LPS+nigericin) condition. The activated condition was set as 0% rescue, and the untreated baseline was considered 100% rescue. The percentage rescue for each treatment condition was determined using the following formula:

[(Activation-treatment)/(activation-baseline)*100], where MCC950 or BFE treatment had experimental values. This calculation facilitated the assessment of how effectively each treatment restored baseline levels of NLRP3 activation by % cells expressing ASC specks and mitochondrial health by ccf-mtDNA % rescue. Statistical comparisons were performed using an unpaired t-test or two-way ANOVA with post-hoc corrections, as appropriate. Comparisons with P<0.05 were considered significant.

### Statistical Analysis

Results were analyzed Microsoft Excel (Version 16.77.1) and GraphPad Prism (version 10.4.1 (532)) for statistical analysis. Data was analyzed using student’s two-tailed *t*-test or an ordinary one-way ANOVA with Tukey or Dunnett’s multiple comparison test to analyze parametric data. The Mann-Whitney U test was used to analyze non-parametric data. To determine if data showed a normal distribution, the Shapiro-Wilk test was used. Outlier analysis was performed using the ROUT method. Values with P<0.05 were considered significant.

## Results

### Clinical, Biological and Social Demographic characteristic of participants

The sample consisted of 10 individuals, 5 healthy controls (CT) and 5 BD patients. Human participant clinical and social demographics (age, gender, and diagnosis) are in presented in Supplementary File 1. Plasma was also extracted from the participant’s whole blood, and ccf-mtDNA was measured and found to be slightly elevated in the plasma of BD patients, in line with previous results seen in the periphery of BD patients^7,32^ (Supplementary Figure 1A). A plasma targeted mitochondrial metabolomic profile (Supplementary Figure 1B) revealed decreased cis-aconitic acid, citric acid, creatinine, and phenylpyruvic acid, alongside increased cystine and nicotinamide adenine dinucleotide (NAD+) - suggesting significant disruptions in energy metabolism, oxidative stress balance, and amino acid metabolism, aligned with previous results^2^.

### Validation and characterization of human derived iPSC and 3D triculture model composed of neural progenitor cells, neurons and astrocytes

All participants recruited for the study (5 BD and 5 CT) underwent PBMC reprogramming into iPSCs. However, only 3 BD-derived and 3 CT-derived iPSC lines passed quality control assessments and were subsequently used for CO generation. Supplementary Figure 2 and Supplementary Table 1 presents the data for the iPSC lines that did not meet quality control criteria. The successful iPSC lines exhibited robust pluripotency marker expression (OCT4, SOX2, TRA-160, ECAD), positive Epi-Pluri-Score (Table 2), and normal karyotyping profiles (Figure 2, Table 3).

**Figure 2.**
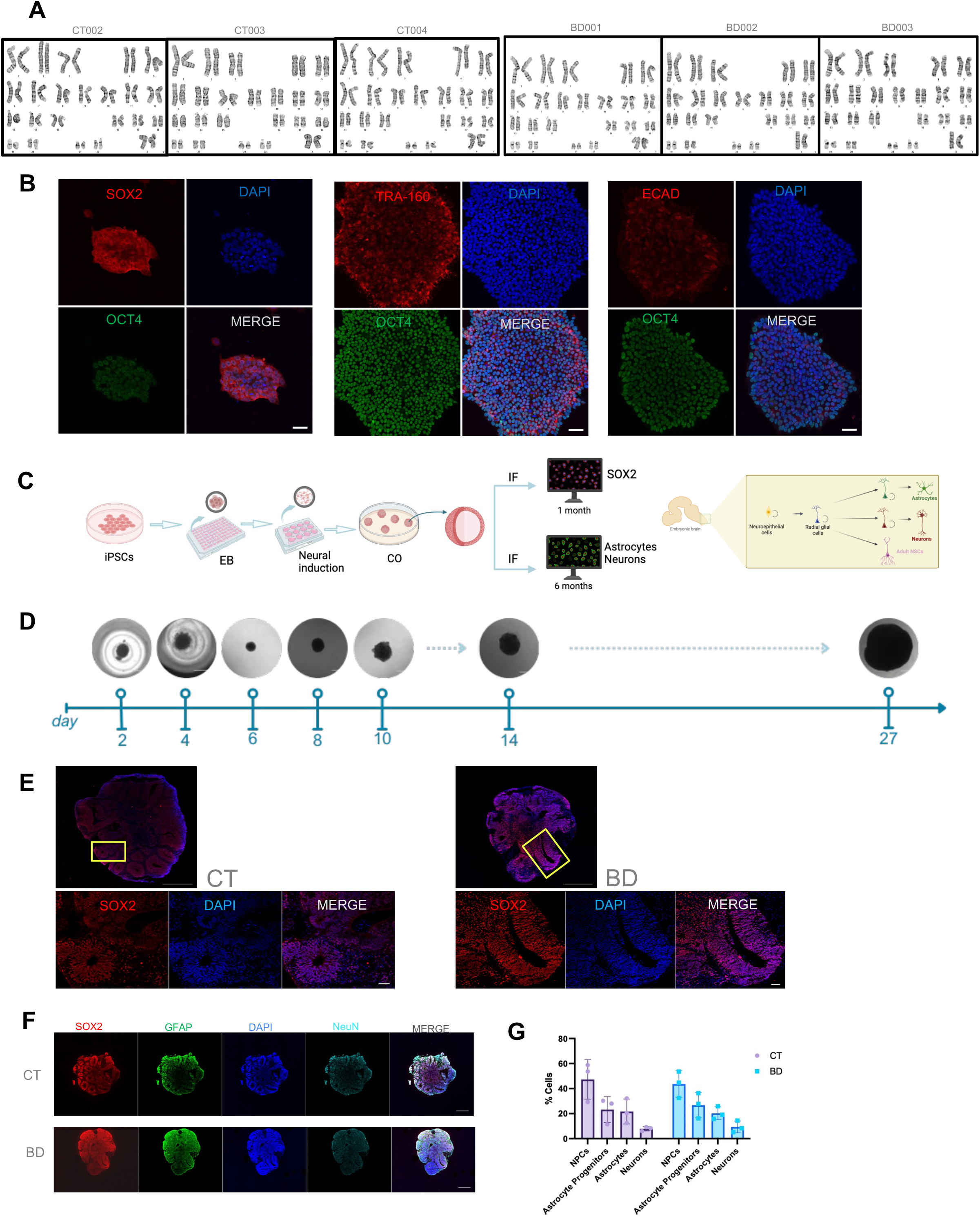
Validation and characterization of iPSC and COs from CT and BD patients. A) Karyotyping analysis of each patient line. B)representative images of immunofluorescence markers for pluripotency SOX2, OCT4, ECAD, TRA-160 in iPSCs. C)Schematic representation of generating COs from iPSCs. D) Representative images of a BD CO at differentiation stages. Scale bar = 1000µm. E) Representative immunohistochemistry images of CT and BD COs at 1 month stained with SOX2 (Red) and DAPI (blue) taken at 10X (whole CO, scale bar=500µm) and 63X (split panels, scale bar = 50µm). F) Representative images of CT and BD COs staining for SOX2(red), GFAP(green), and NeuN(Turquoise), scale bar= 500µm. G) Comparison of cell types across BD and CT COs using HALO image quantification. Bar graphs represent mean ± SD. Data normalized to total cell area.

**Table 3.** Summary of Karyotyping Results.

| Participant # | Karyotyping result | Participant # | Karyotyping result |
| --- | --- | --- | --- |
| CT002 | 46, XX | BD001 | 46, XX |
| CT003 | 46, XX | BD002 | 46, XX |
| CT004 | 46, XX | BD003 | 46, XX |

COs derived from 3 CT and 3 BD patients were grown for 1 month and then underwent characterization to ensure successful differentiation (Figure 2C). The CO generation protocol begins with iPSCs as clumps which form a neuroepithelial boundary and then further differentiate into 3D structures after Matrigel embedding (Figure 2D). In the early stages of CO growth, ventricle-like structures form which contain neural progenitor cells, similar to *in vivo* developmental processes^19,33^. To accurately explore the cytoarchitecture of developing COs, we stained for SOX2 to detect radial glia cells, since most ventricular units display robust SOX2 expression when the CO is 1 month of age and no significant differences were seen across the CT and BD groups for cell types within the COs (Figure 2E-G).

As the CO grows and matures, these SOX2 positive cells further differentiate and give rise to post-mitotic neurons which stain positive for MAP2, and astrocytes which stain positive for GFAP at 6 months (Figure 3B). Together, these markers and developmental timelines demonstrate that the COs generation protocol follows similar patterning and organization to that seen *in vivo*^34^. To gain a deeper understanding of the maturity of COs at 6 months, neuronal local field potential (LFP) recordings revealed distinct activity profiles between the groups. CT COs showed less baseline spike activity when normalized to the maximal CT value. In contrast, BD COs exhibited a significantly higher number of spikes compared to CT COs (Figure 3C). This finding indicates increased neuronal firing in BD COs, suggestive of hyperactive neuronal networks, a characteristic of bipolar disorder reflected in previous literature using animal models of BD^35^. Average spike amplitude measurements showed no significant difference between CT and BD COs (Figure 3D), suggesting that while the frequency of spikes differs markedly between the groups, the intensity of individual spikes remains similar. These findings are supported by work previously done in BD iPSC-derived neurons^36^. The BD COs also exhibited a higher frequency of spikes across various amplitude ranges compared to CT COs, reinforcing the increased spike activity observed in BD samples without a corresponding rise in spike amplitude (Figure 3E, F). This suggests that spikes in BD COs occur more frequently and with greater intensity, reflecting altered neuronal dynamics and potentially prolonged excitatory events. These findings underscore the hyperactive and intense neuronal firing in BD COs, aligning with previously reported neural differences in bipolar disorder^36^.

**Figure 3.**
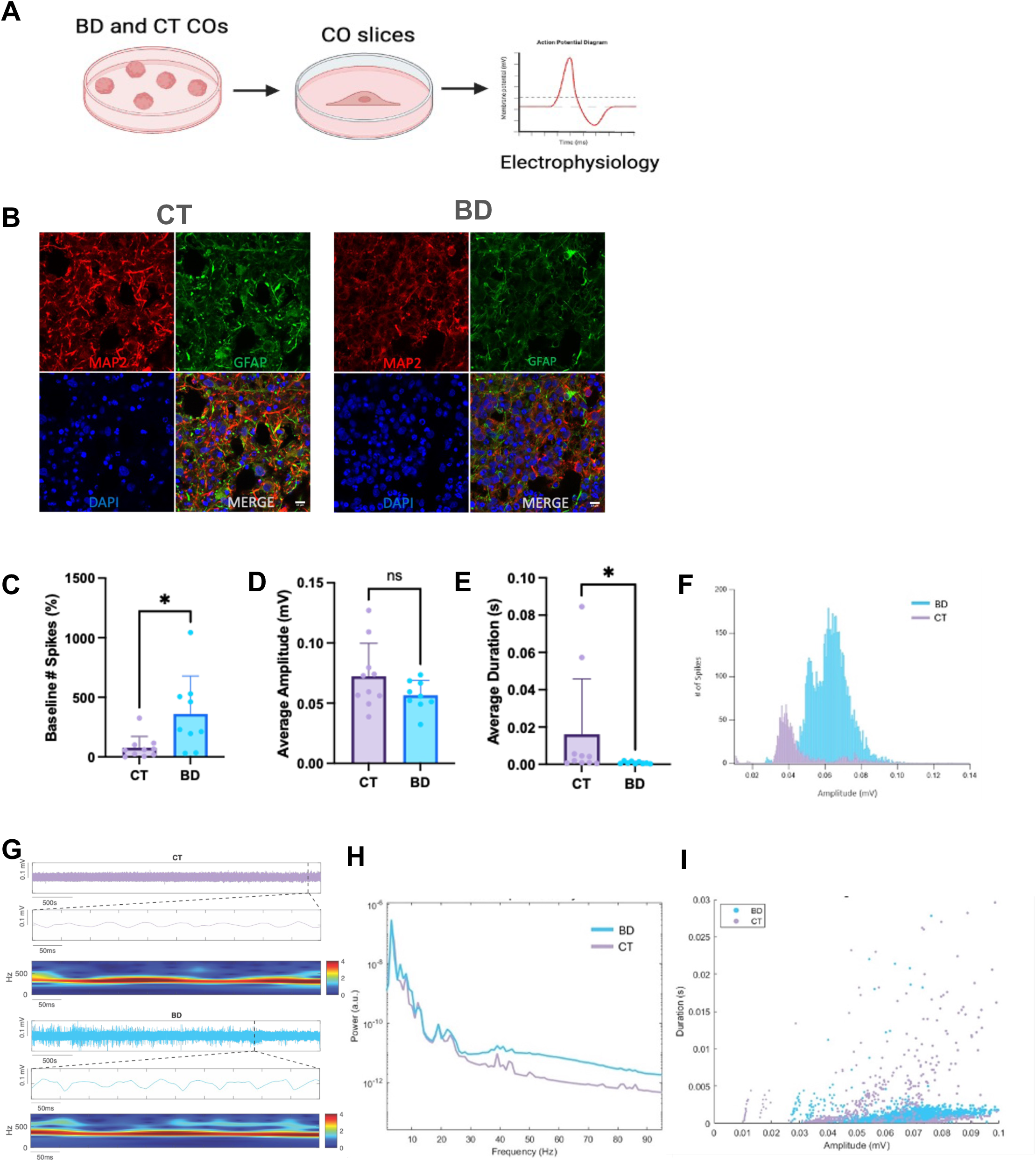
Validation and characterization CO at 6 months maturity. A) Schematic representation of slicing COs to prepare for electrophysiology assessments. B) Representative immunohistochemistry images of CT and BD at 6 months for GFAP (astrocytes) and MAP2 (neurons). Scale bars represent 10µm. C) Normalized to maximal CT, baseline spikes recorded from CT and BD COs. (P=0.013). D)Average spike amplitude in millivolts (mV) from CT and BD COs. E)Average spike duration in seconds. (P=0.0125) (s). F)Histogram plot depicting the relationship between the number of spikes (y-axis) and their amplitude (x-axis). G) Denoised trace representing a 15-minute trace of CT and BD COs and an expanded time segment below each trace with a mortlet plot showing activity between chosen time segments between CT and BD. H) Power spectral density plot. I) Duration scatter plot between CT and BD traces. Data analyzed by Welch’s t-test and Mann-Whitney U test. *P<0.05.

The power spectral density analysis in Figure 3G and 3H revealed a modest increase in power for BD COs relative to CT across most frequencies, with a more pronounced deviation at higher frequencies in BD COs. Scatter plots indicated enhanced intrinsic spiking properties and peak amplitude of local field potentials (LFPs) in BD COs compared to CT, highlighting heightened neuronal activity in the bipolar group (Figure 3I). Collectively, these results suggest a significantly elevated level of neuronal activity in BD COs compared to CT, offering insights into the neural hyperactivity associated with bipolar disorder.

### Reduced Cellular Viability and Altered Metabolomics in BD COs

To gain insight into the intracellular mechanisms altered in BD, the COs at 1 month underwent extensive metabolic profiling (Figure 4A). Previous studies have noted that BD COs are of a smaller size compared to healthy controls^37^. Interestingly, when the COs were dissociated to single cells, the BD COs were found to have significantly fewer cells per CO compared to the CT (Figure 4B). COs then underwent mitochondrial metabolomic profiling, normalized to cell number, using 20 selected metabolites based on previous work from our group investigating metabolic syndrome in BD with a much larger patient cohort^2^. Notably, the BD COs trended lower in ATP and glucose-6-phosphate levels (Figure 4C). This could explain peripheral metabolic dysregulation, specifically, impaired whole body glucose homeostasis in often seen in BD patients^38^. These findings also align with previous studies showing that BD patients experience widespread dysregulation in energy metabolism, particularly affecting downstream ATP-dependent processes^39^. To further confirm these results, live single cells from COs were dissociated to measure intracellular ATP using Cell Titer Glo (Figure 4D). BD COs again showed a significant reduction in ATP levels, further emphasizing the metabolic vulnerabilities associated with BD .

**Figure 4.**
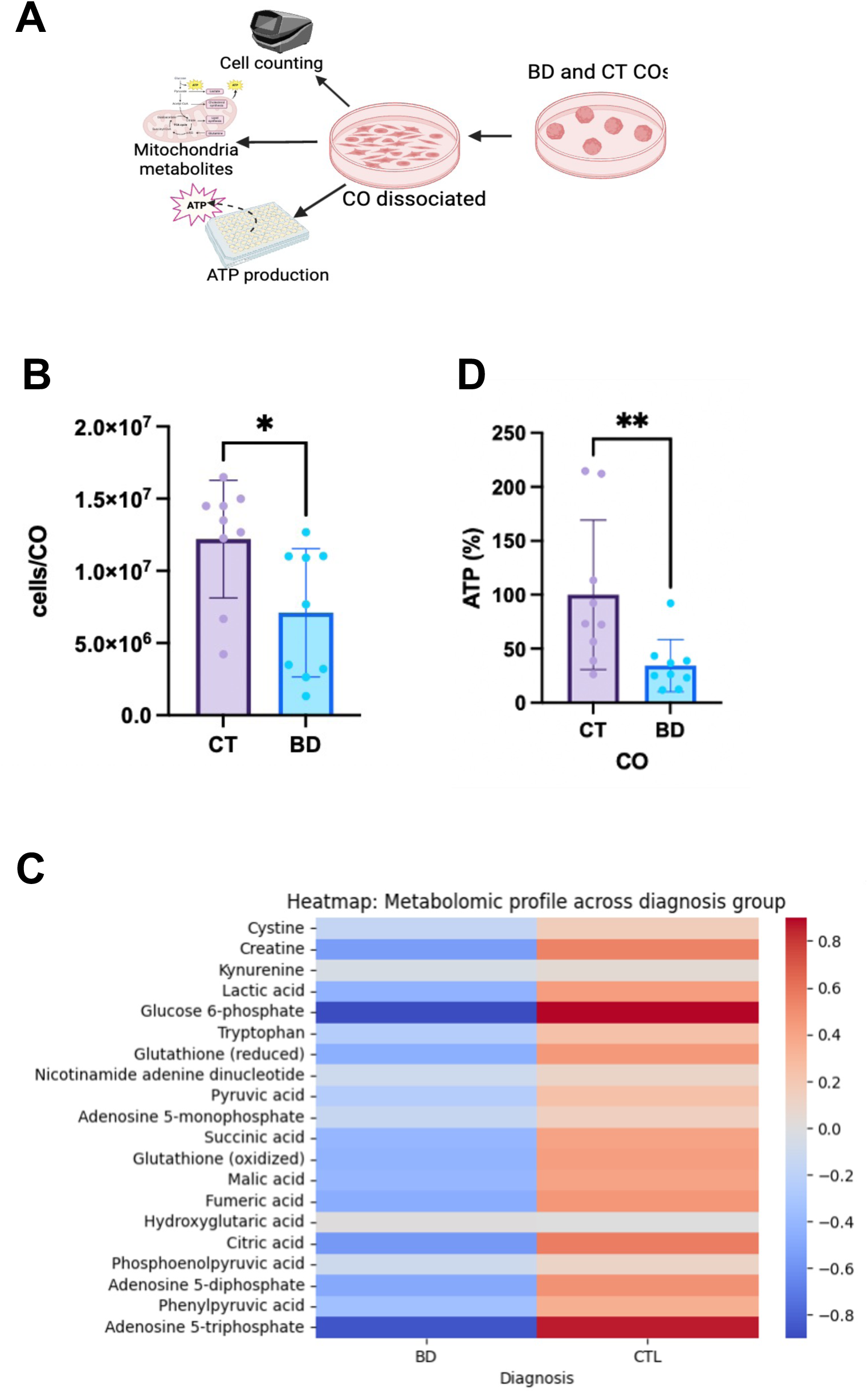
Reduced Cellular Viability and Altered Metabolomics in BD Cerebral Organoids. A) Schematic representing the experiments done on the COs for bioenergetic and electrophysiology assessments. B) cell number in each CO was measured. Each data point indicates 1 CO dissociated. Data analyzed by Mann-Whitney U. (P=0.0082). C) Heatmap of intracellular mitochondrial metabolites stratified by BD and CT CO groups. D) ATP levels across BD and CT Groups. Data analyzed by Mann-Whitney U. (P=0.004). Bar graphs represent mean ± SD. *P<0.05; ** P<0.005.

### Mitochondrial Health in BD COs: Altered Mitochondrial Membrane Potential, Dynamics, and Morphology

Altered energy metabolism is strongly associated with abnormal mitochondrial morphology and function. Therefore, mitochondrial function and morphology in COs was measured^40^. Using transmission electron microscopy (TEM), numbers of round (immature) versus elongated (mature) mitochondria were measured. While both groups had similar levels of elongated mitochondria, the BD group had a significantly higher number of round mitochondria, consistent with previous literature on post-mortem brains of BD patients^40^ (Figure 5B, 5C). Next, overall mitochondrial mass was examined using MitoTracker Green, a fluorescent compound which accumulates in the mitochondrial matrix to label mitochondrial mass in the cells^41^. Images were taken using a confocal microscope, and upon quantification, similar levels of mitochondrial mass in the BD and CT COs were found (Figure 5D, 5E).

**Figure 5.**
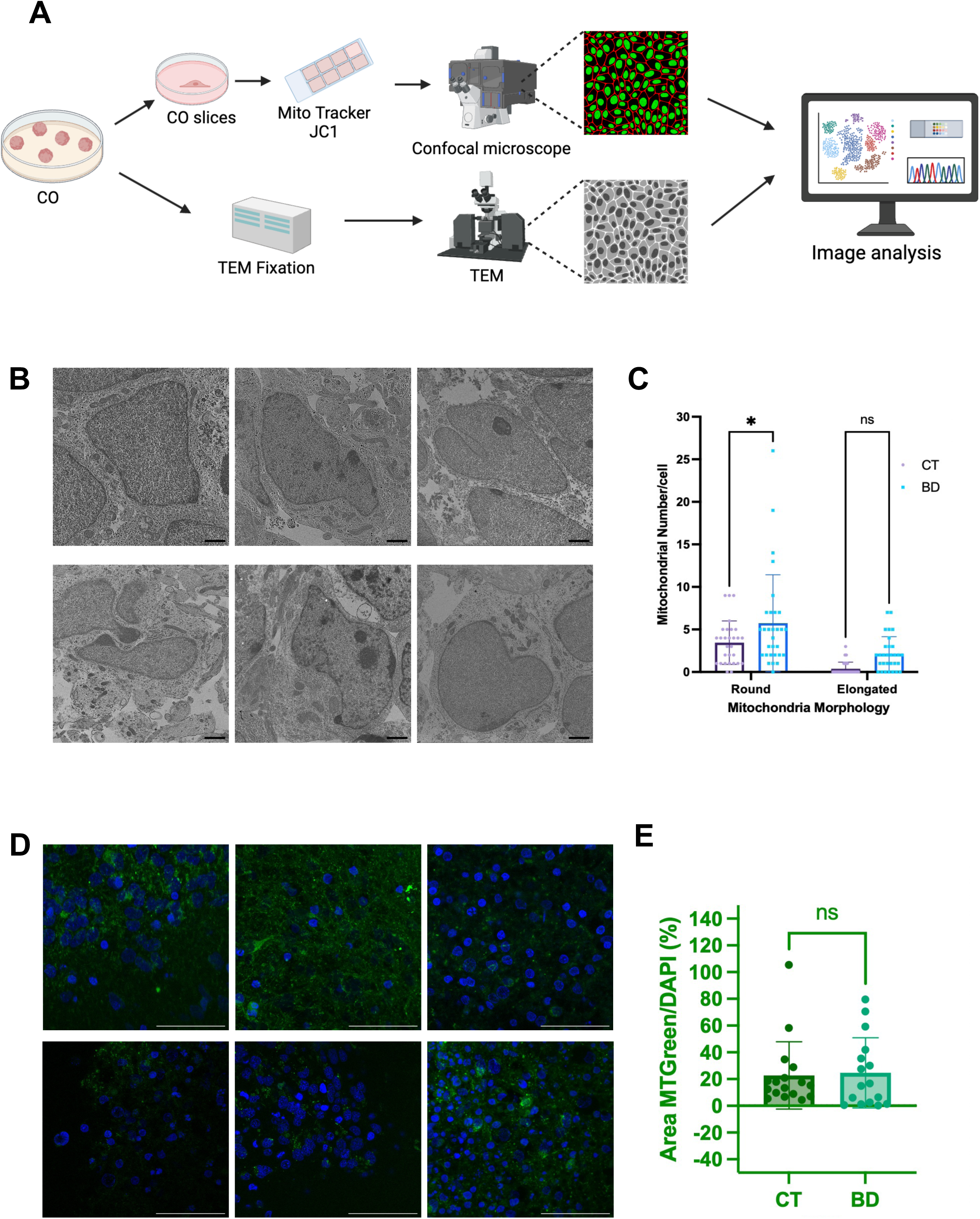
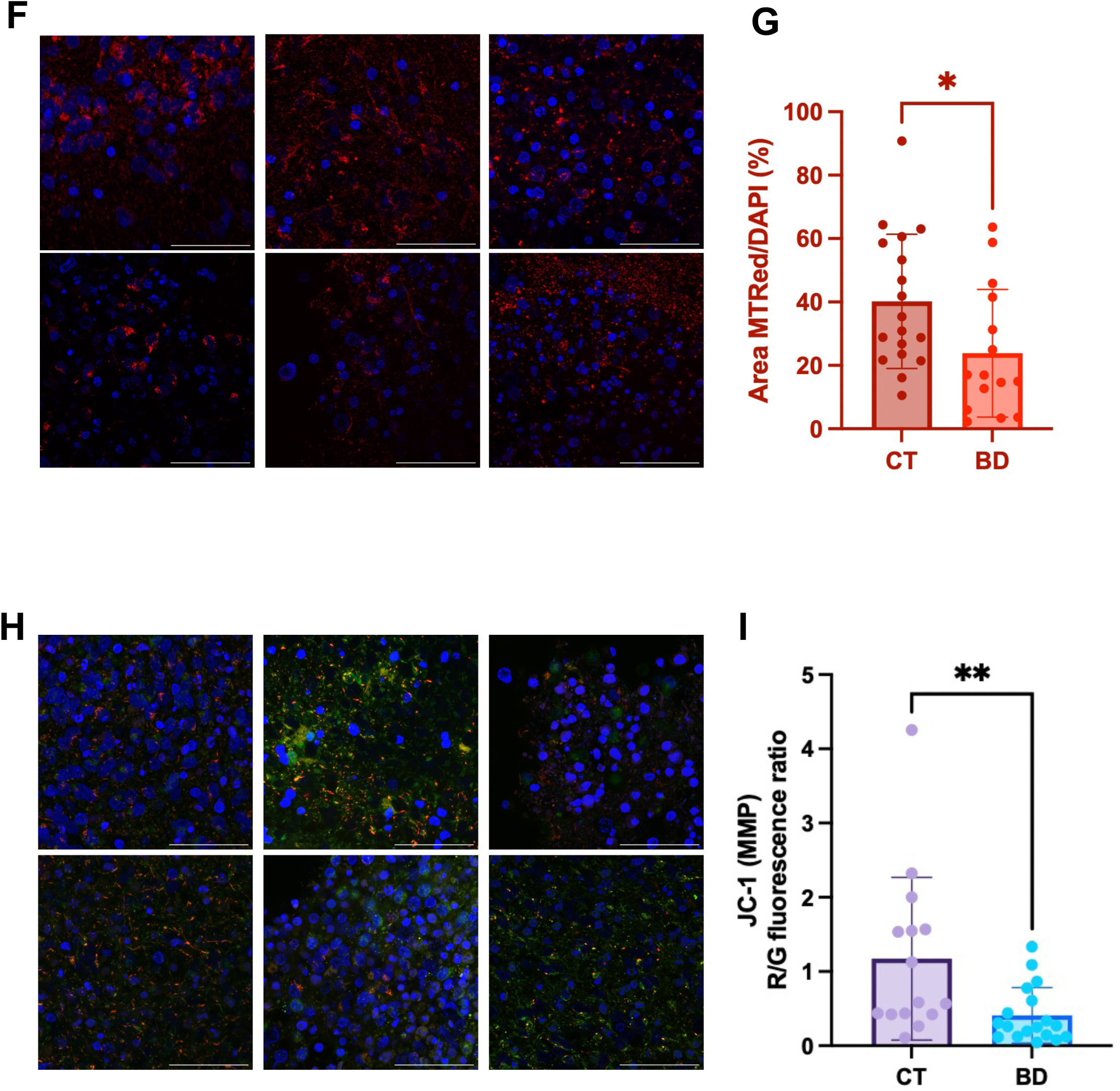
Mitochondrial Dysfunction in BD Cerebral Organoids: Altered Mitochondrial Membrane Potential, Polarity, and Morphology. A) Schematic representing experimental protocol to examine mitochondrial dysfunction in COs. B) TEM images scale bar = 5µm. C)) quantifying number of round and elongated mitochondria and data analyzed by two-way ANOVA. (P=0.018) for round, (P=0.087) for elongated. D-E) Representative images of MitoTracker Green and quantified using HALO and analyzed by Mann-Whitney U test. Scale bar = 50µm. F-G) Representative images of MitoTracker Red CMX ROS and quantified using HALO and analyzed Welch’s t-test. (P=0.031). Scale bar = 50µm H-I) Representative images of red to green ratio using JC-1 fluorescent dye. Scale bar = 50µm and images quantified using HALO and analyzed by Mann-Whitney U test. (P=0.005). Bar graphs represent mean ± SD. *P<0.05; **P<0.005.

While there were no significant differences in mitochondrial mass, the remaining question was whether this translates to healthy, functional mitochondria. The first approach was to stain the COs using MitoTracker Red CMX ROS, a cationic fluorescent dye which solely penetrates live, healthy mitochondria in a potential-dependent manner^41^ (Figure 5F). After quantifying the images, the mitochondria in the CT COs stained at a significantly higher intensity with MitoTracker Red than those in the BD group, suggesting that the mitochondria are in better health, and more hyperpolarized compared to the mitochondria in the BD group (Figure 5G). JC-1, a dual emission cationic dye was used to indirectly evaluate changes in the mitochondrial membrane potential (MMP) (Fig 5H). This dye enters the cells and accumulates in the mitochondria in response to their MMP, yielding a red fluorescence for highly energetic mitochondria and green fluorescence for weaker mitochondria^42^. Image quantification revealed that the BD COs exhibited a significantly lower MMP in comparison to the CT (Figure 5I), corresponding to a weaker energetic state as suggested by the MitoTracker Red stain results. Together, these findings highlight the relationship between morphological changes in the mitochondria of BD COs and their ability to maintain a polarized, functional membrane potential compared to CT.

### Enhanced Extracellular Markers of Inflammation and Mitochondrial Stress in BD COs

Given the increased evidence of mitochondrial dysfunction seen in the BD COs, along with increased evidence of oxidative stress in BD, we next sought to explore the extracellular status of the COs^43^. Mitochondrial dysfunction within the cell often leads to overproduction of reactive oxygen species (ROS), which can subsequently trigger the release of DNA into the cytosol^44^. To assess this, we measured ROS levels in the supernatant and found significantly elevated levels in BD COs compared to CT COs (Figure 6B). Additionally, BD COs showed significantly higher levels of extracellular double-stranded DNA (dsDNA) (Figure 6C).

**Figure 6.**
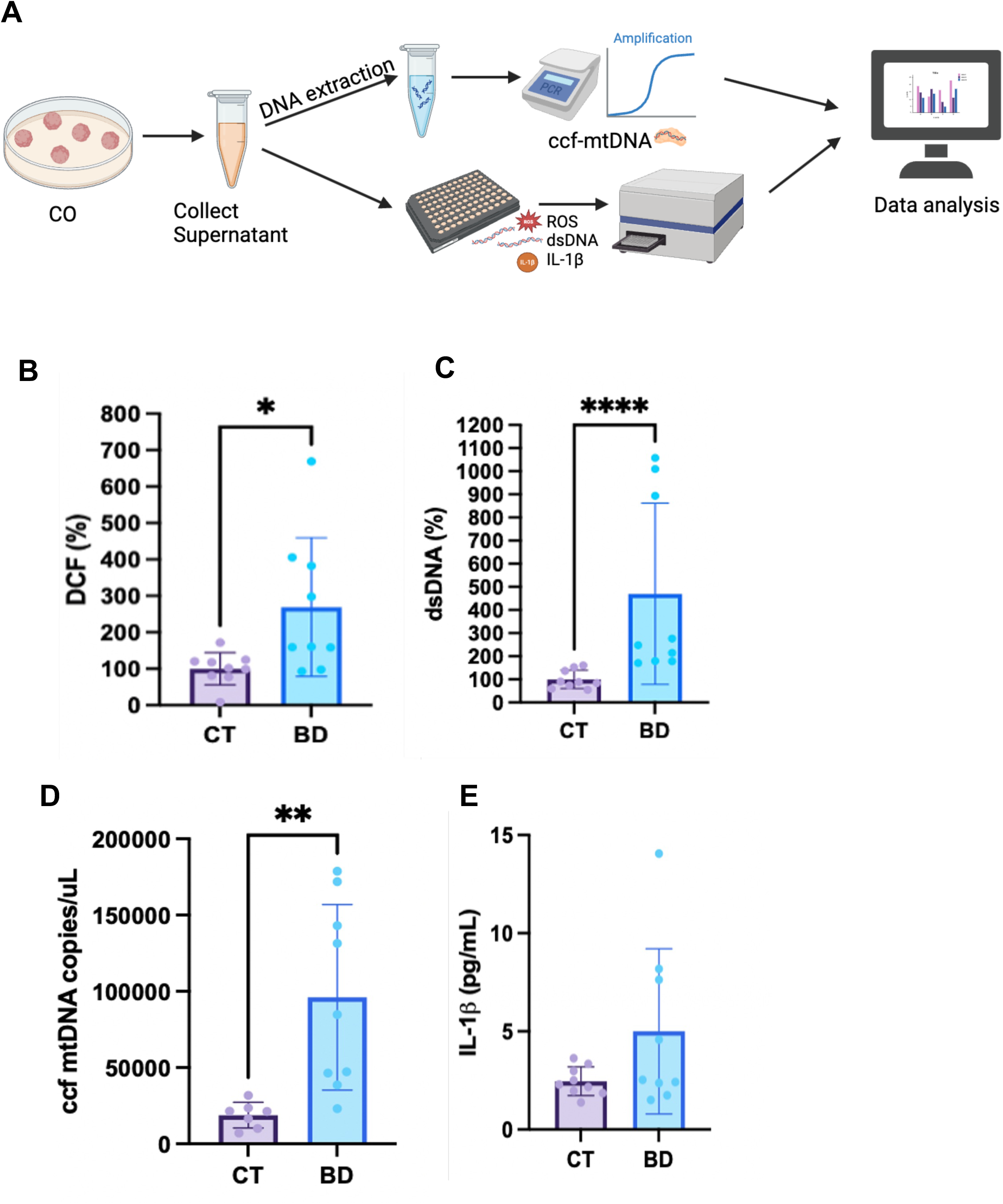
Enhanced Extracellular Markers of Inflammation and Mitochondrial Stress in BD Cerebral Organoids. A) Schematic representing experimental protocol to measure extracellular markers of CO stress. B) measurement of ROS. Data analyzed by Welch’s T-test. (P=0.029) (DCF) levels in COs. Data analyzed by C) measurement of dsDNA release. Data analyzed by Welch’s T-test. P=0.04. D) measurement of ccf-mtDNA. Data analyzed by Welch’s T-test. (P=0.005). E) measurement of il-1β levels. Data analyzed by Mann-Whitney U. (P=0.026). Bar graphs represent mean ± SD. *P<0.05; **P<0.005; ****P<)0.00005

Upon increased stress, mitochondria can release their DNA through mitochondrial permeability transition pore opening^2^; we specifically measured ccf-mtDNA to further investigate this phenomenon. BD CO media contained significantly higher levels of ccf-mtDNA, indicating much of the double stranded DNA released is coming from the mitochondria (Figure 6D). Inflammatory markers in the periphery and in post-mortem brain samples of BD patients have been increasingly reported in the literature, suggesting there is an underlying phenomenon of neuronal inflammation^45,46^. To assess the downstream inflammatory response, we measured levels of IL-1β as a marker of inflammation. Although the difference was not statistically significant, BD COs exhibited a trend toward higher IL-1β levels compared to CT (Figure 6E).

### Increased Susceptibility to NLRP3 Inflammasome Activation in BD COs

Previous studies have shown that elevated ROS and ccf-mtDNA can influence NLRP3 activation both directly and indirectly^44^. Our group has also previously shown in human blood that patients with mood disorders are more sensitive to NRLP3 activation^14^, and developed an appropriate technical approach to detect NLRP3 activation in 2D and 3D tissue using LPS, a primer, and Nigericin, an NLRP3 inflammasome activator^14,47^. The NLRP3 inflammasome is a tripartite complex composed of NLRP3, ASC, and pro-caspase-1. Upon activation, the complex assembles and promotes inflammation which can be visualized through formation of ASC specks^14^. The NLRP3 inflammasome is present in both astrocytes and microglia in the human brain, and both cell types have been used together or separately to study NRLP3 activation^48^.

The COs generated come from an ectodermal differentiation lineage, containing neurons and astrocytes, not microglia^47^. Hence, we assessed NLRP3 activation in astrocytes of the COs both at baseline and following NLRP3 activation to evaluate the sensitivity of BD COs to inflammasome injury compared to CT COs. Figure 7 illustrates the heightened vulnerability of BD COs to NLRP3 inflammasome activation at baseline and after induced injury using LPS and Nigericin. Representative immunofluorescence images of ASC specks (marking activated NLRP3), demonstrate a 141% increase in NLRP3 activation in the CT and a 275.8% increase in activation in the BD group (P=0.0479). This resulted in a 1.9 fold increase in activation sensitivity in BD COs compared to CT, suggesting the BD CO’s are more prone to NLRP3 activation (Figure 7A, B). Upon activation, levels of ccf-mtDNA were also measured and found to be significantly elevated in both BD and CT groups (Figure 7C).

**Figure 7.**
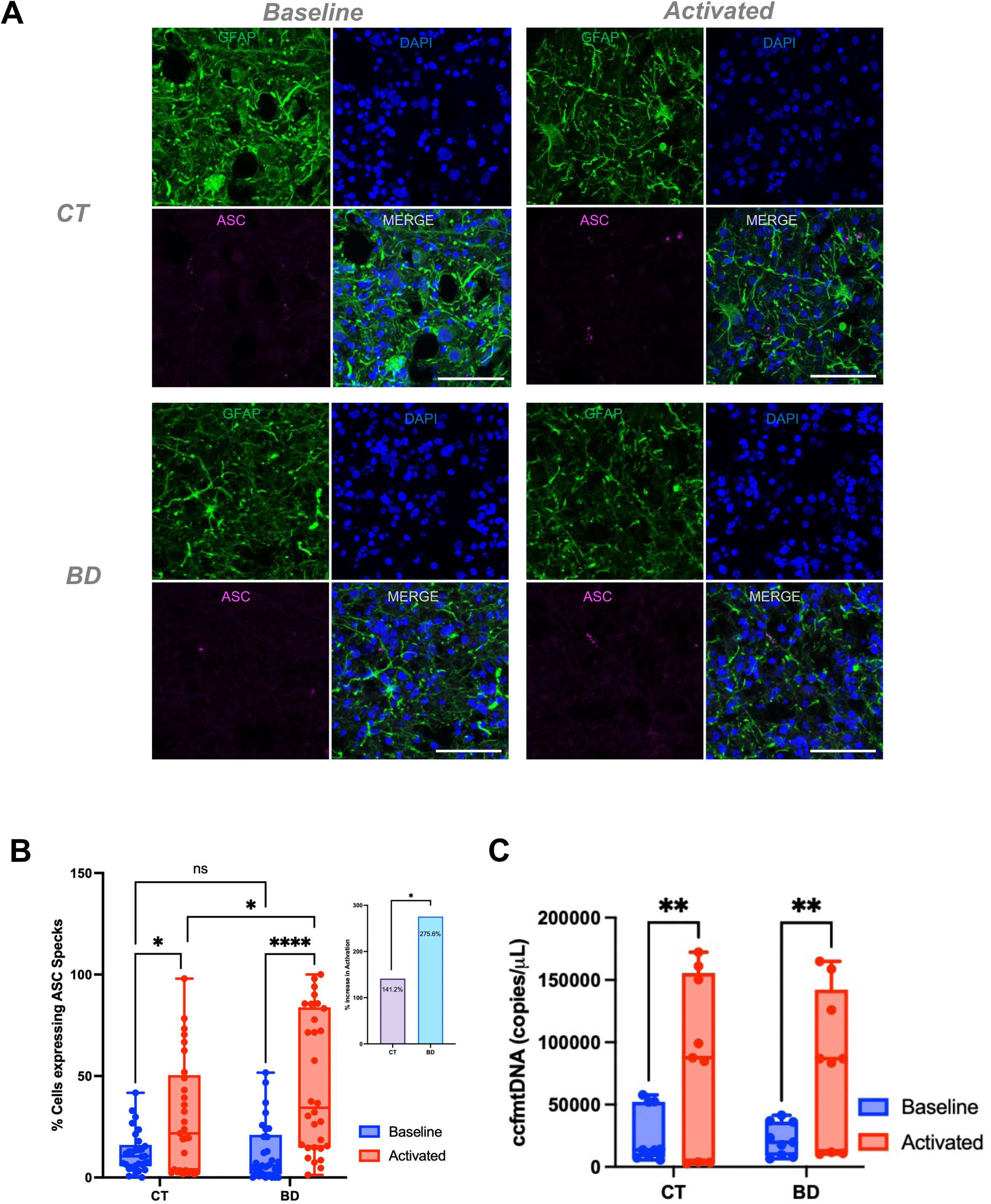
Increased Susceptibility to NLRP3 Inflammasome Activation in BD Cerebral Organoids. A) Representative images staining GFAP for astrocytes (green), DAPI (blue), and ASC (magenta). Baseline represents untreated control, activated is LPS priming for 3 hours, followed by nigericin for 4 hours. Scale bar =50µm B) quantification of ASC specks using HALO and ImageJ as described in the methods.((Baseline CT vs Activated CT P=0.027); (Baseline BD vs Activated BD P<0.0001); (Activated CT vs Activated BD P=0.0479)). C)levels of ccf-mtDNA upon activation. P<0.005. Data analyzed by Two-way ANOVA Tukey’s multiple comparison’s test. Bar graphs represent mean ± SD. *P<0.05; **P<0.005; ****P<)0.00005

### Using a novel Bioactive Flavonoid Extract to rescue NLRP3 activation

We decided to explore targeting the NLRP3 inflammasome with an inhibitor to attempt to rescue the increased susceptibility of BD COs. Currently, MCC950 is the leading compound used to inhibit the NLRP3 activation, however it has been implicated with hepatotoxicity ^49,50^. Our group has previously published on the use of a BFE, derived from *Euterpe oleracea Mart.* (açaí), which has been extensively studied by Machado group and has consistently demonstrated antioxidant and anti-inflammatory effect via partial inhibition of the NLRP3 inflammasome activation^28,29,31,51^.

To assess the CO’s therapeutic potential of BFE treatment, COs were pre-treated with BFE or MCC950 (Figure 8A). ASC specks were quantified, and we found decreased specks in both MCC950 and BFE treatment conditions (Figure 8B). To determine the efficacy of MCC950 and BFE in modulating NLRP3 activation, we quantified the specks and calculated their rescue effects relative to the activated condition (Figure 8C). MCC950 treatment led to a rescue of 113.2% in CT COs and 125.3% in BD COs, indicating that it effectively restored activation levels beyond baseline in both groups, with a slightly greater effect in BD. The lower rescue effect of BFE in BD COs may be attributed to the higher initial activation levels in BD compared to CT, making complete restoration to baseline more challenging. This finding suggests that the exaggerated inflammasome activation in BD COs may limit the efficacy of certain modulators, requiring further exploration into whether higher doses or combinatorial treatments could enhance the rescue effect in BD.

**Figure 8.**
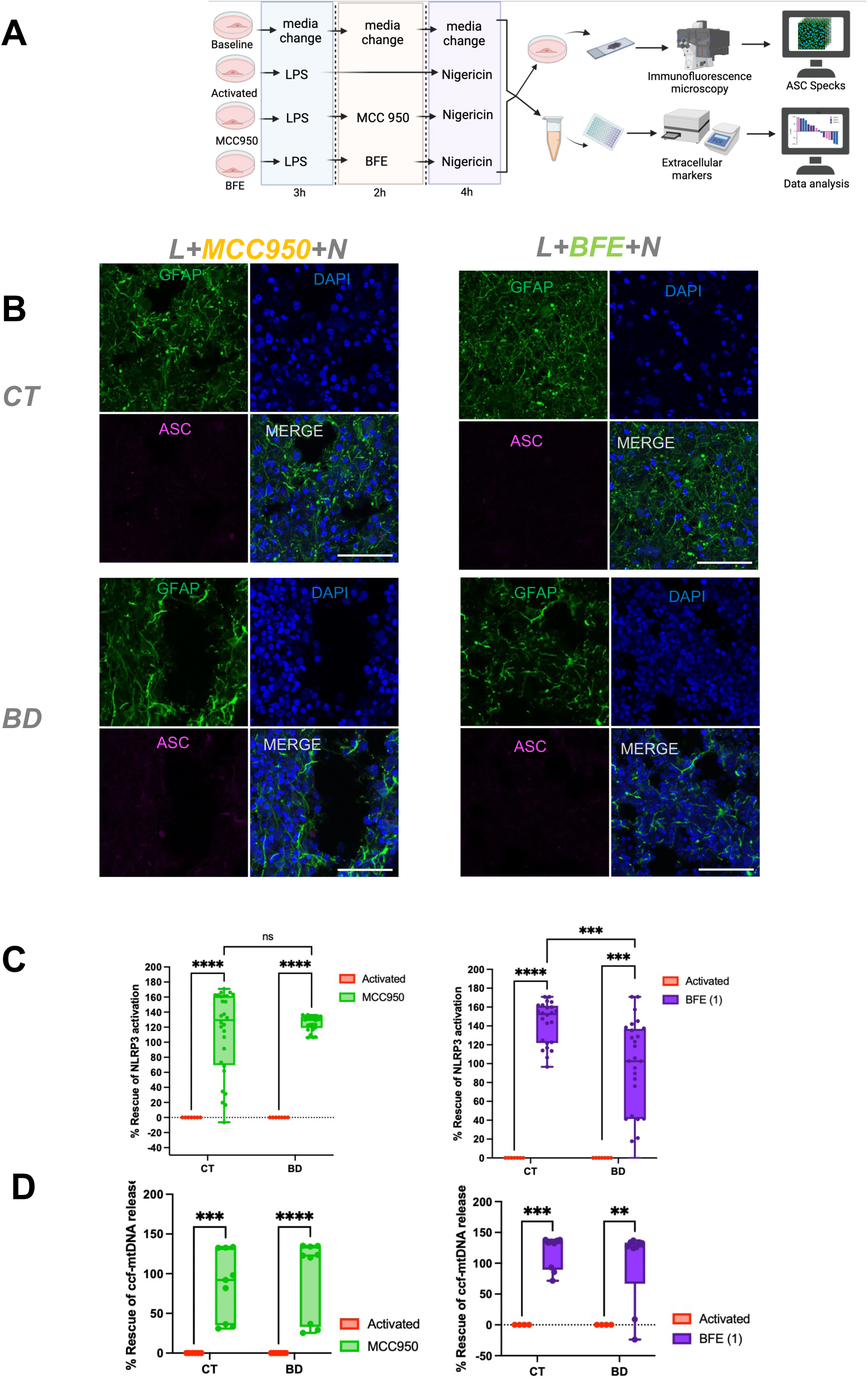
BFE rescues injury of NLRP3 inflammasome in COs. A)Schematic representing experimental method to activate NLRP3 inflammasome in COs. B) Representative images staining GFAP for astrocytes (green), DAPI (blue), and ASC (magenta) Scale bar =50µm. Slices were primed with LPS for 3 hours, followed by 2 hours of MCC950 or BFE, followed by nigericin for 4 hours. C) quantification of ASC specks using HALO and ImageJ as described in the methods, presented as a % rescue of NLRP3 activation. Bar graph represents mean ± SD. ((Activated CT vs MCC950 CT P<0.0001); (Activated BD vs MCC950 BD P<0.0001); (Activated CT vs BFE CT P<0.0001); (Activated BD vs BFE BD P=0.0009)). D) measurement of %rescue of ccf-mtDNA release. ((Activated CT vs MCC950 CT P=0.0006); (Activated BD vs MCC950 BD P<0.0001); (Activated CT vs BFE CT P=0.0004); (Activated BD vs BFE BD P=0.0024)). Bar graph represents mean ± SD. Data analyzed by Two-way ANOVA Tukey’s multiple comparison’s test. *P<0.05; **P<0.005; ****P<)0.00005

Alongside NLRP3 activation, circulating cell-free mitochondrial DNA (ccf-mtDNA) release was also significantly reduced following MCC950 and BFE treatment, indicating that both interventions mitigated mitochondrial stress in BD and CT COs (Figure 8D). These findings reinforce the link between mitochondrial dysfunction and inflammasome activation, and the variation in rescue magnitude between CT and BD suggests underlying differences in inflammasome and redox regulation and mitochondrial dysfunction between the two groups, warranting further investigation into their distinct inflammatory and mitochondrial profiles.

## Discussion

BD is a complex neuropsychiatric mood disorder with several interconnected theories about its etiology. To date, no existing study mechanism has solely demonstrated sufficient accuracy in depicting disease etiology. Therefore, our objective was to explore the interplay between two key players in BD disease pathophysiology: mitochondrial dysfunction and inflammation. Our use of patient-derived COs allowed us to investigate disease specific phenotypes which are too complex to model in an animal. Unlike traditional 2D cultures, the 3D COs partially mimic the organization and patterning seen *in vivo*, including interactions between neurons and astrocytes in a physiologically relevant environment.

Accumulating evidence from structural and functional MRI studies have shown abnormal size and impaired function in the frontal and prefrontal cortex of BD Patients^52,53^. Interestingly, *in vitro* studies using COs have reported smaller sphere sizes in the BD compared to the CT groups^37,54^. It was no surprise to find that the BD COs generated by our group had a reduced cell number compared to CT. Various hallmarks of energy metabolism have also been found in specific brain regions and blood platelets of BD patients where evidence suggests aberrations in mitochondrial bioenergetics as well as energy production^53^. When examining metabolomic profiling in the COs, the BD COs seemed to have compromised glucose metabolism and energy production. These metabolic alterations can contribute to energy production and oxidative stress. Several studies have found evidence of impaired ATP production in BD. Konradi et al. (2004) reported a pronounced decrease (78%) in expressions of genes which regulate oxidative phosphorylation and ATP-dependant processes in BD^39^. Similarly, Andreazza et al. (2010) observed a significant decrease (∼53%) in mitochondrial complex I activity in the brain of BD subjects, which could lead to reduced ATP production^8^. These studies collectively support our results which show depleted ATP production in the BD COs.

Ion pump dysfunction, altered calcium homeostasis, and irregular action potential generation are all downstream effects of depleted ATP and are documented in BD^55^. Previous research has demonstrated the detectability of neuronal activity in human-derived COs through various electrophysiological and imaging methodologies^56–59^. When we examined LFP recordings of 6-month-old CT and BD patient-derived COs, we observed an increase in baseline spiking activity in BD COs compared to CT. The significant increase in spike frequency in BD COs indicates an overall increase in neuronal firing rate, consistent with the hyperactivity often seen in BD animal models^35^. The elevated baseline activity observed in BD COs also aligns with previous studies on 2D neuronal cultures from bipolar patients that showed hyperexcitability^36,60^. While the neuropathological differences in BD have been investigated in some of the characteristics of the medial prefrontal cortical network and anterior cortex the overall connectivity and network require further investigation^61,62^.

Recently, impairment of mitochondrial energy metabolism has gained great attention in BD pathophysiology^55^. Post-mortem brain staining of BD prefrontal samples have revealed morphological abnormalities and abnormal distribution of mitochondria^63,64^. In our COs, using transmission electron microscopy, we found BD COs had a significantly higher number of round mitochondria compared to CT. Round mitochondria are often associated with oxidative stress, uncoupling, and mitochondrial fragmentation, while elongated mitochondria are more common in healthy cells and are associated with enhanced OXPHOS^64,65^. This finding is consistent with a study in neurons derived from BD iPSCs which showed significantly smaller mitochondrial size compared to CT^36^. Some studies have shown that round mitochondria can lower MMP^66^. This was seen in our BD Cos, which had a significantly lower MMP and lower quantities of polarized mitochondria. All the above results indicate a strong connection between perturbed mitochondrial function and BD.

Mitochondrial dysfunction also contributes to decreased mitochondrial immunity through elevated levels of ROS and ccf-mtDNA^44^. Decreased MMP and altered mitochondrial bioenergetics contribute to weakened mitochondrial integrity which causes mtDNA release into the periphery, referred to as ccf-mtDNA. Increased oxidative stress has been consistently reported in both post-mortem brain samples and blood samples of patients with BD^8,9,17^. These high levels of ROS, when accumulated, can cause functional impairments in cells. BD COs showed increased ROS, ccf-mtDNA, and dsDNA. These findings provide a comprehensive view of the extent of dysregulated cellular homeostasis in BD Cos^15^.

High ccf-mtDNA and ROS has also been widely associated with poor mitochondrial health and greater symptom severity in patients with BD^7,9,10,32^. The biological role of ccf-mtDNA and mtDNA fragments has been associated as a potential link between the brain and immune system^67^. Ccf-mtDNA can act as a DAMP, suggesting that inflammation is secondary to a primary intrinsic process caused by compromised mitochondria^67^. In healthy cells, mtDNA is rapidly removed from the cytosol through nucleases. However, in diseased states, or during infection, the amount of DNA released from a cell can overwhelm the capacity of these nucleases^15,17,68^. Ccf-mtDNA in the cytosol can also activate toll-like-receptors (TLRs), upregulate interferons, and promote transcription of NLRP3 genes^69,70^. Our group has previously reported data on 837 BD patients suggesting ccf-mtDNA as a marker of chronic low-grade inflammation^2^. We have also previously reported increased protein expression of NLRP3 in post-mortem brain tissue of BD patients and that PBMCs of patients with mood disorders can be increasingly sensitive to NLRP3 activation, possibly due to underlying inflammation^16,71^. Therefore, this prompted us to investigate NLRP3 activation sensitive in the BD COs.

Astrocytes are the most abundant type of glial cells found in the brain with various physiological roles to maintain brain function and immunity^72^. Consistent with findings reported in the literature, the BD COs showed increased sensitivity to NLRP3 activation in the astrocytes. Recent studies have shown astrocytes to be potential contributors to inflammatory abnormalities sin mood disorders. Specifically, inflammatory upregulation in astrocytes has also been associated with reduced brain volume and cognitive impairment in schizoprenia^73^. This highlights the potential role of NLRP3 inflammasome as a link between mitochondrial dysfunction and inflammation^74^. Mitochondria possess a pivotal role in both neuron and astrocyte function in the brain. They regulate energy production, neuronal firing, and neuroprotection. Mitochondrial impairments in mood disorders are highly associated with neuroinflammatory states^72^. The loss of MMP, elevated ROS, and ccf-mtDNA in the BD COs made them more prone to NLRP3 activation, which, in turn, further increased ccf-mtDNA therefore, causing the COs’ homeostasis to be disrupted both intracellularly and extracellularly.

Several major neuropsychiatric diseases including mood disorders and BD exhibit some form of mitochondrial dysfunction and heightened inflammation, yet treating each separately has not yet been shown to be effective. The NLRP3 inflammasome integrates both aspects of mitochondrial stress and inflammatory triggers, making it a desirable target. The NLRP3 inflammasome is also involved in other neurodegenerative diseases. Parkinson’s disease (PD) is a neurodegenerative disease with strong links to mitochondrial dysfunction and NLRP3-drive inflammation. Interestingly, NLRP3 deficient PD murine models showed protection from PD, making it a desirable therapeutic target^15,75^.

Progress in targeted treatment for mood disorders has been limited, mostly due to the incomplete understanding of the biological mechanisms underlying the onset and severity of the disease. MCC950 is a classical small molecule NLRP3 inhibitor that can block assembly of the NLRP3 inflammasome oligomerization complex^76^. This drug has been widely considered as an anti-inflammatory alternative to treat NLRP3 induced inflammation, including in autoinflammatory and autoimmune diseases^77^. However growing concerns over hepatotoxicity of MCC950 in phase II clinical trials underlines the need for alternatives^49,50^.

To further investigate the vulnerability of the BD COs, an attempt was made to rescue the COs using a well-established anti-inflammatory antioxidant extract, BFE. Our group has conducted extensive research to develop the BFE extract. BFE is a hydrolyzed purified chemical extract from a native berry in the Brazilian Amazon rainforest with numerous bioactive properties, which act synerigistically^31^. It is an antioxidant, neuroprotective, and anti-inflammatory agent composed of several potent naturally-occurring flavonoids including catechin and epicatechin^31^. BFE has consistently demonstrated anti-inflammatory potential, and successful modulation of the NLRP3 inflammasome by our group using microglia, VERO cells, and THP-1 derived macrophages^28,31,78^. In a 2021 study, BFE demonstrated a reduction in the inflammatory response triggered by olanzapine, an antipsychotic drug in macrophages^51^. This suggests BFE can be used as a potential agent to treat or prevent neuroinflammation in neuropsychiatric diseases. In our most recent *in silico* study using BFE in Davidson et al. (2024), molecular docking and molecular dynamic simulations revealed that the main bioactive molecules in BFE diminished inflammatory response driven by LPS+nigericin NLRP3 activation^27^. Decreased NLRP3 activation would therefore be predicted through pre-treatment with BFE, consistent with our findings in the COs.

### Limitations

While this study revealed novel findings, it is important to acknowledge the limited sample size (3 BD and 3 CT). Hence, these results can only guide our knowledge of BD but not be used to make strong conclusions about the disease. Additionally, an ongoing challenge in generating COs is maturation. COs do not contain any vasculature; thus, the core of the COs can become necrotic as inadequate oxygen, and nutrients are able to diffuse into the core^19^. The COs we currently use are a simplified model of the fetal brain, containing cells that primarily come from ectodermal differentiation. However, they do not fully recapitulate the entire human brain as they lack other components such as microglia, which can further help us understand the neuroinflammatory effects in BD. While it may be possible to generate microglia-containing COs, these spontaneous formation, self-directed protocols are not yet robust and well established in terms of cell ratio and heterogeneity between patient iPSC lines. Additionally, co-culturing microglia with COs can also hinder the physiological relevance of the COs as they are now being matured in a different environment which does not fully recapitulate microglial physiological activity. However, with recent scientific and technological advances, we can begin generating more complex models and versions of COs containing vasculature and increased cell types. This in turn will help us increase utilization of COs for investigating the origins of mitochondrial dysfunction in neurological diseases and metabolic health.

## Conclusions

Multiple psychiatric disorders stem from one commonality: cerebral metabolic dysregulation, and it is the mitochondria which plays a pivotal role in these pathways^79^. To examine how mitochondria may drive inflammatory and neuropsychiatric diseases, it is important to first understand how mitochondria are behaving under normal healthy cell signalling. Mitochondrial biogenesis is a critical process to maintain cellular energy homeostasis to allow cells to easily adapt to the body’s varying energy demands^79^. The results in the BD COs suggest that mitochondria are unable to meet the CO’s metabolic demands which further explains why we see altered metabolomics as well as aberrant electrophysiological activity. This metabolic deficit was accompanied by increased neuronal firing rates, suggesting a connection between mitochondrial dysfunction and hyperactive neuronal networks, a hallmark of BD. Intriguingly, overlapping pathways, if disrupted in key brain regions which control metabolism, could also explain why individuals with severe mental illness, including BD, experience metabolic dysfunction, even pre-morbidly^80,81^.

While it has been widely known that the mitochondria are the ATP machines of the cell, they can also act like reservoirs for DAMPs. Our group has previously published on a cohort of youth BD showing high levels of mtDNA in serum were significantly associated with more severe depression symptoms and higher lactate, another indicator of mitochondrial dysfunction^7^. Elevated levels of ccf-mtDNA have been reported in BD patients^2,32^. This currently is also evident in our findings when looking at ccf-mtDNA levels in BD patient plasma as well as BD COs. In our study, ccf-mtDNA is possibly being the crosslink between mitochondrial impairment and the NLRP3 activation. BD COs were significantly more sensitive to NLRP3 inflammasome activation, responding with heightened inflammatory reactivity compared to CT. Notably, this is the first experimental study to conduct an analysis involving NLRP3, COs, and BD, providing a foundational framework for future investigations.

Importantly, we also provide insight into novel therapeutic strategies, demonstrating that targeting NLRP3 inflammasome inhibition using BFE effectively mitigates inflammatory responses and restores cellular homeostasis. These findings emphasize the need for integrated therapeutic approaches that simultaneously address mitochondrial dysfunction and inflammation, offering new avenues for intervention in BD.

## Acknowledgements

We would like to thank AM’s group in Brazil for their expertise in formulating BFE D.E.S.E.S. was supported by a MITO2i Graduate Scholarship for this work. All figures were *Created in BioRender. Zachos, K. (2025)* https://BioRender.com/ *c51y326*.

