## Supplementary Figure for "iPSC-Derived Cerebral Organoids Reveal Mitochondrial, Inflammatory and Neuronal Vulnerabilities in Bipolar Disorder"

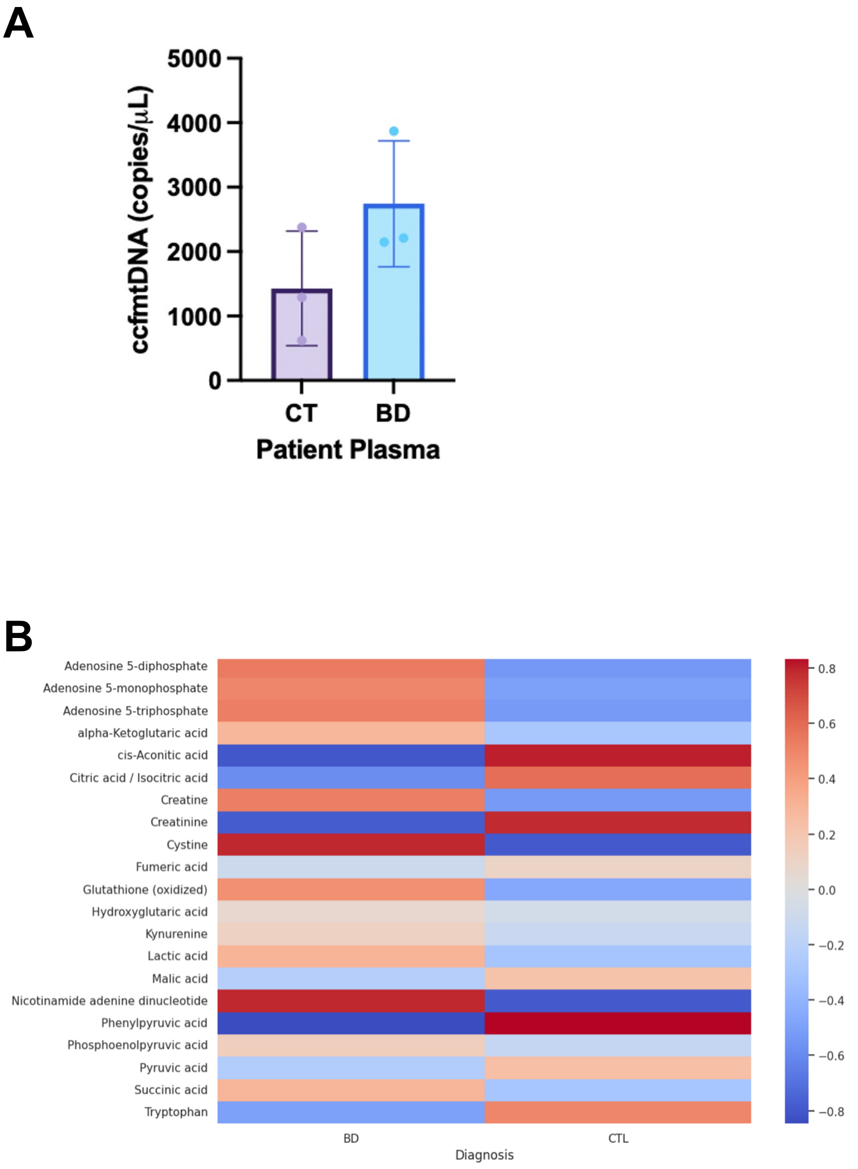


**Supplementary Figure 1.** Data collected from patient plasma. A) measurement of ccf mtDNA (copies/µL) Data analyzed by Welch's t-test. P=0.16. B) Heatmap of mitochondrial plasma metabolites stratified by disease.

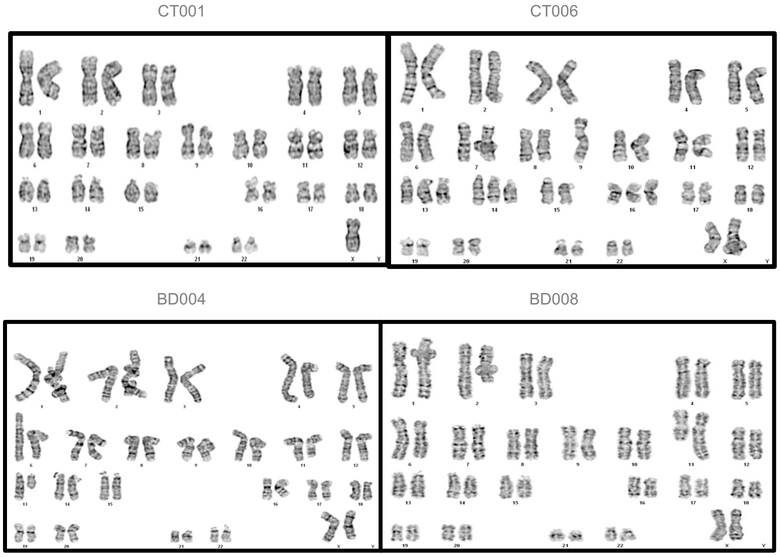


**Supplementary Figure 2**. Abnormal karyotyping of additional participants

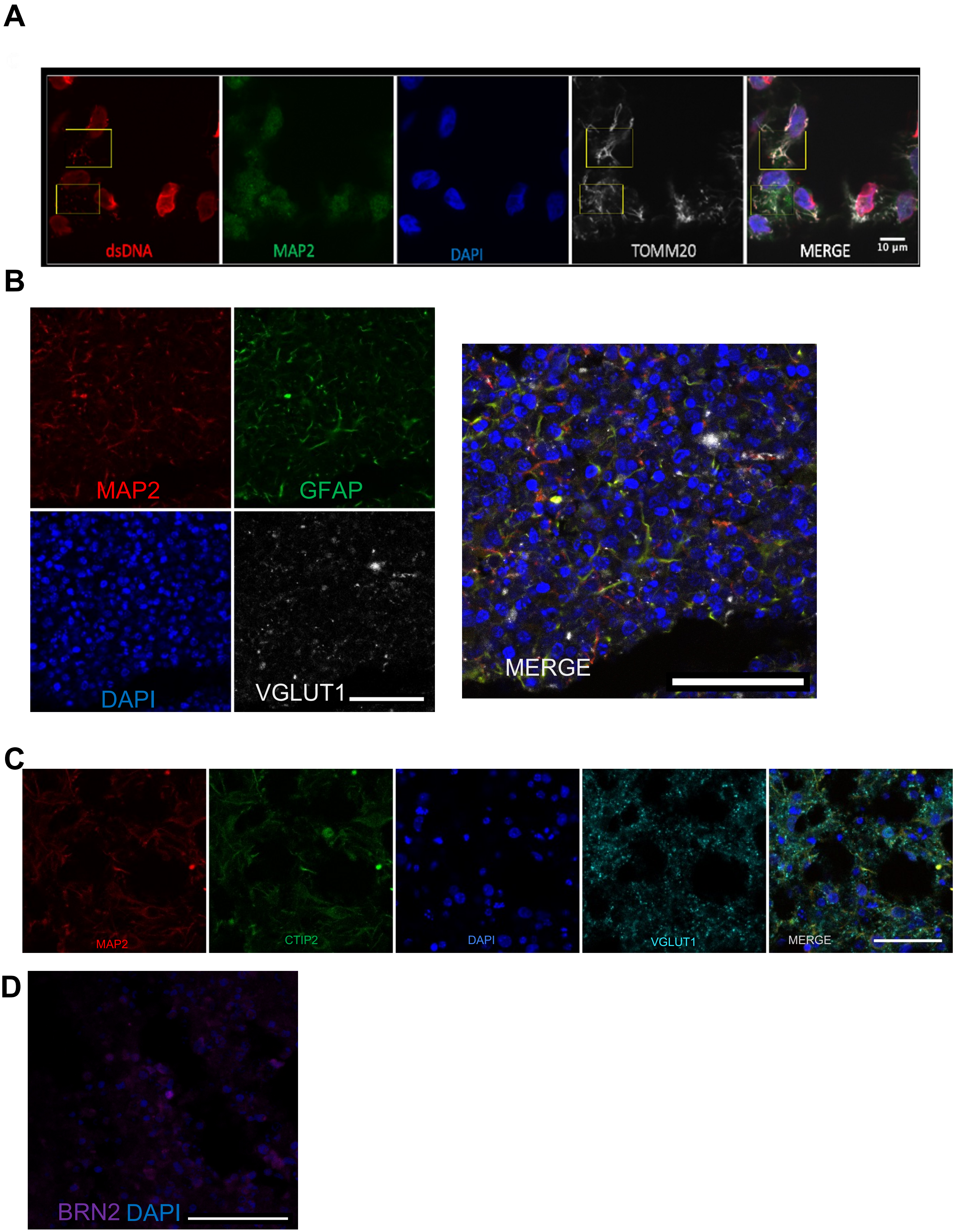


**Supplementary Figure 3.** Additional CO staining. A) TOMM20, dsDNA, MAP2 markers in 1 month old COs. Scale bar = 10um B) MAP2, GFAP, VGLUT1 in 6 month old COs. Scale bar = 50um. C) BRN2 in 6 month old COs. D) MAP2, VGLUT1, CTIP2 in 6 month old COs. Scale bar = 50um

**Supplementary Table 1.** Summary results table of karyotyping

| Participant # | Karyotyping result | Participant # | Karyotyping result |
|---|---|---|---|
| CT001 | Mosaic, 46, X, -X,+mar/46,XX[20] | BD004 | Abnormal, 46XX, add(11)(q23)[cp6] |
| CT006 | Mosaic, 46-48, XX, add(9)(q?13)[cp2]/46, XX[16] | BD008 | Mosaic, 46, XX, t(2;11)(q2?3;q23)[2]/46,XX[5] |
